# Multiple host targets of *Pseudomonas* effector protein HopM1 form a protein complex regulating apoplastic immunity and water homeostasis

**DOI:** 10.1101/2023.07.31.551310

**Authors:** Kinya Nomura, Lori Alice Imboden, Hirokazu Tanaka, Sheng Yang He

**Author notes:** Corresponding author: Sheng Yang He Department of Biology Duke University Durham, NC 27708.

## Abstract

Bacterial type III effector proteins injected into the host cell play a critical role in mediating bacterial interactions with plant and animal hosts. Notably, some bacterial effectors are reported to target sequence-unrelated host proteins with unknown functional relationships. The *Pseudomonas syringae* effector HopM1 is such an example; it interacts with and/or degrades several HopM1-interacting (MIN) Arabidopsis proteins, including HopM1-interacting protein 2 (MIN2/RAD23), HopM1-interacting protein 7 (MIN7/BIG5), HopM1-interacting protein 10 (MIN10/14-3-3ĸ), and HopM1-interacting protein 13 (MIN13/BIG2). In this study, we purified the MIN7 complex formed *in planta* and found that it contains MIN7, MIN10, MIN13, as well as a tetratricopeptide repeat protein named HLB1. Mutational analysis showed that, like MIN7, HLB1 is required for pathogen-associated molecular pattern (PAMP)-, effector-, and benzothiadiazole (BTH)-triggered immunity. HLB1 is recruited to the trans-Golgi network (TGN)/early endosome (EE) in a MIN7-dependent manner. Both *min7* and *hlb1* mutant leaves contained elevated water content in the leaf apoplast and artificial water infiltration into the leaf apoplast was sufficient to phenocopy immune-suppressing phenotype of HopM1. These results suggest that multiple HopM1-targeted MIN proteins form a protein complex with a dual role in modulating water level and immunity in the apoplast, which provides an explanation for the dual phenotypes of HopM1 during bacterial pathogenesis.

## Introduction

The type III protein secretion system (T3SS) is a major virulence factor in many Gram-negative bacterial pathogens of plant and animals. This secretion system translocates a large number of effector proteins into the host cell to promote pathogenesis. *P. syringae* pv. *tomato* (*Pst*) strain DC3000 is a bacterial pathogen of tomato and Arabidopsis and, in the past three decades, has been used as a model to understand bacterial pathogenesis and the virulence functions of type III effectors in plants [1]. *Pst* DC3000 produces about 36 effectors [2]. The host targets of many effectors from *Pst* DC3000 and related *P. syringae* strains have been reported. Identification of these host targets gives a glimpse into an impressive array of infection-associated cellular processes, ranging from pattern-triggered immunity to apoplast water homeostasis, that are manipulated by bacteria during pathogenesis [1, 3-8].

HopM1 and AvrE are two of the most highly conserved type III effectors in *P. syringae* [9]. These two effectors share no sequence similarity, but are functionally redundant; they collectively make a large virulence contribution to *Pst* DC3000 infection in susceptible plants Arabidopsis, tomato and *Nicotiana benthamiana* [10-13]. Bacterial delivery of HopM1 is associated with suppression of defense-associated callose deposition in the plant cell wall [12, 14], suppression of vascular flow [15], and induction of “water-soaking” in the apoplast of infected leaves [16-18]. In addition, transgenic overexpression of HopM1 results in suppression of apoplastic oxidative burst and stomatal closure [19]. In a previous study, we identified several Hop<u>M</u>1-<u>in</u>teracting (MIN) Arabidopsis proteins, including MIN2 (RAD23a), MIN7 (an ARF-GEF) and MIN10 (14-3-3 KAPPA) [20]. Importantly, the *min7* knockout mutant plants could partially rescue the virulence defect of the *Pst* DC3000 *hopM1 avrE* double mutant (the ΔEM mutant hereinafter) or the ΔCEL mutant, in which several <u>C</u>onserved-<u>E</u>ffector-<u>L</u>ocus-associated effector genes, including *hopM1* and *avrE* are deleted [20, 21]. [Note: because the virulence phenotypes of the ΔCEL mutant and the ΔEM mutant are essentially identical [21], we used these strains interchangeably in this study.] The partial restoration of ΔEM mutant and ΔCEL mutant multiplication in the *min7* knockout mutant plants suggests that, in the absence of a biologically relevant host target (e.g., MIN7), HopM1 is no longer fully necessary for the bacterium to cause disease [20]. Both HopM1 and MIN7 were found to be localized to the trans-Golgi network (TGN)/early endosome compartment (EE), suggesting HopM1 attacks a vesicle trafficking pathway as part of its virulence mechanism to promote bacterial virulence [22, 23].

In contrast to the *min7* mutant, other *min* mutant plants did not rescue the virulence defect of the *Pst* DC3000 ΔCEL mutant. However, most *MIN* genes belong to multi-gene families. For example, *MIN10* belongs to the 14-3-3 protein family, which has 14 members in Arabidopsis [24]. Therefore, a lack of disease phenotype in other *min* single mutant may be because of functional redundancy among members of these particular gene families. Indeed, it was shown that chemicals known to inhibit client protein interaction with 14-3-3 proteins could rescue the virulence defect of the ΔCEL bacterial mutant [25], suggesting that targeting 14-3-3 proteins, as a group, may be biologically relevant to the HopM1 function. Still, why HopM1 is able to target multiple sequence-unrelated host proteins and functional relationships between HopM1-targeted host proteins remain enigmatic. Similarly, why HopM1 affects both immune response and water homeostasis in the leaf apoplast is not clear. Furthermore, HopM1 was found to also interact with several E3 ubiquitin ligases and proteasome subunits [26] and to activate autophagy [27] and an Arabidopsis mutant defective in plasm membrane-localized abscisic acid (ABA) transporter ABCG40 can also genetically restore the virulence of the ΔEM mutant in Arabidopsis leaves [18]. It is currently not known whether diverse host proteins (e.g., MIN proteins, E3 ubiquitin ligases, proteasome subunits and ABA transporters) associated with HopM1 virulence functions are involved in related or separate processes and whether they are involved in the virulence function of HopM1 at different stages and/or different host cell types during bacterial infection.

Besides HopM1, other *P. syringae* effectors have been reported to target more than one host protein. For example, AvrPto and AvrPtoB target a number of immune-associated transmembrane receptor-like kinases, as well as the salicylic acid receptor NPR1 protein [28-36]. AvrPphB cleaves several cytoplasmic/membrane-associated kinases including PBS1, BIK1, PBS1-like proteins and CPK3 [36, 37]. HopAI1 has been shown to interact and inactivate mitogen-activated protein kinases MPK3, MPK4, and MPK6 [38, 39]. In these cases, most of the host targets for each effector are related in sequence. Accordingly, the biological relevance of a candidate host target(s) to the function of the cognate effector during bacterial infection is likely dependent on its relative binding affinity and subcellular location *in vivo*. For several other effectors, the identified host targets are unrelated in sequence. For example, AvrB has been shown to interact with RIN4 (a novel protein), MPK4 (a mitogen-activated kinase), RAR1 (a CHORD-domain protein) and HSP90 (a heat shock protein) [40, 41]. HopF2 targets RIN4, BAK1 (a transmembrane receptor-like kinase) and MKK5 (a mitogen-activated kinase kinase) [42-45]. HopZ1a acetylates HID1 (2-hydroxyisoflavanone dehydratase) in soybean and tubulin, MKK7 JAZ transcriptional repressors in Arabidopsis [46-48]. Why a single effector could target multiple, sequence-unrelated host proteins is a significant puzzle in the study of pathogen effectors and this puzzle is likely to become more prominent as more pathogen effectors in bacteria, fungi and oomycetes are subjected to detailed investigations.

In this study, we discovered that MIN7 exists in a ∼700 kDa protein complex in Arabidopsis leaf cells. This unexpected finding prompted us to perform *in vivo* protein pull down and co-immunoprecipitation (co-IP) experiments to characterize the MIN7 complex. These experiments led to the realization that MIN7, MIN10 and MIN13 are components of the MIN7 complex *in vivo*. Furthermore, we identified additional components of the MIN7 complex, including 14-3-3 CHI (related to MIN10), ARFA1c (a substrate of ARF-GEF) and a tetratricopeptide repeat (TPR) protein, HLB1 (HYPERSENSITIVE TO LATRUNCULIN B1; [49]). Analysis of knockout mutant plants showed that, like the *min7* mutant, the *hlb1* mutant is compromised in multiple branches of immunity. HLB1 is localized to the TGN/ EE compartment in a MIN7-dependent manner. In addition, like the *min7* mutant, the *hlb1* mutants contain an elevated water content in the apoplast and, notably, artificial water supplementation to the leaf apoplast was sufficient to phenocopy callose-suppressing phenotype of HopM1.

Taken together, these results provide an explanation for the ability of HopM1 to interact with sequence-unrelated host MIN proteins. Multiple HopM1-targeted host MIN proteins appear to form a protein complex required for the proper interplay between water homeostasis and immunity in the leaf apoplast, providing an explanation for the ability of HopM1 to perturb water homeostasis, immune responses and possibly other cellular processes in the leaf apoplast.

## Results

### Isolation of the MIN7 Protein Complex *in vivo*

The first indication that the MIN7 protein might be in a protein complex was from native polyacrylamide gel electrophoresis (PAGE) analysis of the MIN7 protein in detergent-solubilized Arabidopsis TGN/EE vesicle preparations (see Materials and Methods). The predicted molecular weight (MW) of MIN7 is ∼200 kDa (1,739 aa). However, the apparent MW of MIN7 in the native PAGE gel was ∼700 kDa (Fig. 1A), suggesting that MIN7 is in a large protein complex in Arabidopsis leaf cells.

**Figure 1.**
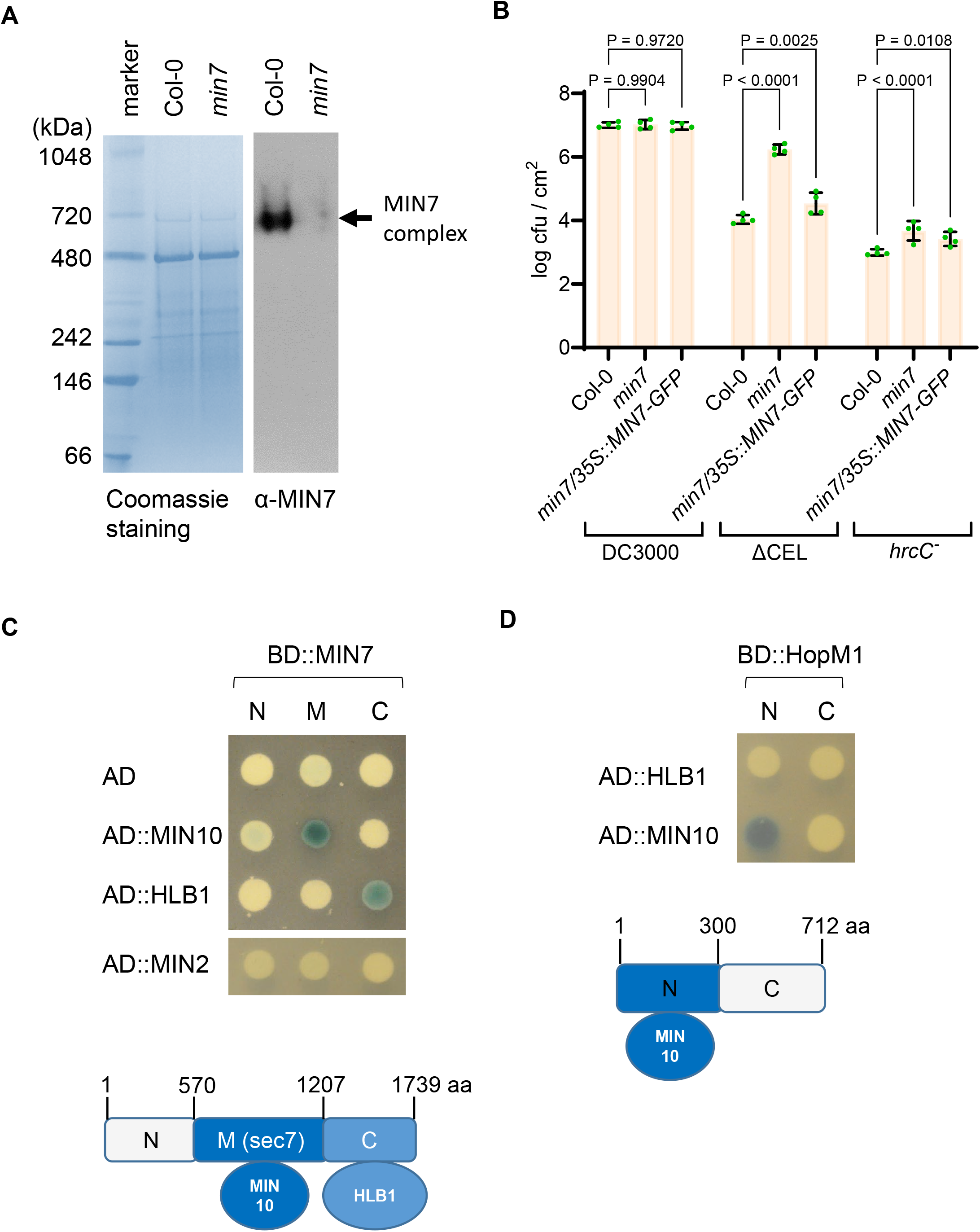
The MIN7 protein complex in Arabidopsis. (A) Detection of MIN7 complex by native PAGE analysis. Solubilized TGN protein extract containing MIN7 (∼200kDa) was separated by native PAGE. MIN7 complex (∼700 kDa) was detected by MIN7-specific antibody. (B) Bacterial multiplication in Col-0, *min7* and *min7*/*35S::MIN7-GFP* plants. Arabidopsis plants were dip-inoculated with *Pst* DC3000, the ΔCEL mutant or the *hrcC* mutant (at 1×10^8^ cfu/ml) and immediately covered with a clear plastic dome to maintain high humidity. Bacterial populations (mean ± SEM; n=4 leaf samples) in leaves were determined at day 4 post dip-inoculation. (C) Yeast two-hybrid (Y2H) interaction assay. Truncated MIN7 proteins (-N, -M, and -C) were expressed from pGILDA (BD fusion). MIN2, MIN10 and HLB11 were expressed from pB42AD (AD fusion). Yeast cultures were spotted and grown on minimal medium containing galactose and X-gal. Blue color indicates interaction, whereas white color indicates no interaction. (D) Y2H interaction assay between HopM1 and HLB1. Truncated HopM1 (N and C) were expressed from pGILDA (BD fusion). HLB1 and MIN10 were expressed from pB42AD (AD fusion). Yeast cultures were spotted and grown on minimal medium containing galactose and X-gal. Blue color indicates interaction, whereas white color indicates no interaction.

To further test the hypothesis that MIN7 is in a protein complex and to identify components of the MIN7 complex, we performed GFP-Trap® A-based protein pull down experiments. For this purpose, we first transformed the Arabidopsis *min7* mutant with a 35S:MIN7-GFP construct, which expresses a MIN7-GFP fusion protein under the control of the CaMV 35S promoter. Independent lines expressing various levels of MIN7-GFP were obtained and propagated to T2 and T3 generations. Line 1 expresses MIN7-GFP at a level comparable to the endogenous MIN7 and was used for further analysis. Most importantly, we determined the functionality of the MIN7-GFP fusion protein by assessing the ability of the MIN7-GFP fusion to complement the *min7* mutant. As reported before, the ΔCEL mutant multiplied poorly in wild-type Col-0 plants, but could multiply aggressively in the *min7* mutant plants (Fig. 1B). Transgenic expression of MIN-GFP restored Arabidopsis resistance to the ΔCEL mutant (Fig. 1B), confirming that MIN7-GFP is functional.

GFP-Trap® A-based protein pull down experiments were performed three times with detergent-solubilized total protein extracts from MIN7-GFP plants and, as controls, extracts from Col-0 and Col-0/35S::GFP plants. The protein compositions were analyzed by mass spectrometry (see Materials and Methods). Proteins found in MIN7-GFP pull down, but not in Col-0 or Col-0/35S::GFP (S1 Table) pull down, are listed in Table 1. Interestingly, we detected not only MIN7, but also MIN10 and MIN13. MIN13 was not described in the study of Nomura et al. (2006) due to its relatively weak, but detectable yeast two-hybrid interaction with HopM1 (Fig. S1). In addition to MIN10 and MIN13, several proteins that were not identified as HopM1 interactors in our previous study were also found, including 14-3-3 CHI (related to MIN10), ARFA1c (a substrate of ARF-GEF) and HLB1 [24, 49, 50] (Fig. S2). Üstün and colleagues (2014) previously identified several E3 ubiquitin ligases and proteasome subunits, in addition to MIN7 protein, in HopM1 pull down experiments in the *N. benthamiana* transient expression system. These *N. benthamiana* HopM1-interacting proteins were not found in the Arabidopsis MIN7 complex in the absence of HopM1.

**Table 1.** Arabidopsis proteins pulled down with MIN7-GFP *in vivo^a^*.

| Identified Proteins | Accession # | #1 | #2 | #3 |
| --- | --- | --- | --- | --- |
| **Pulled down from detergent-solubilized total leaf extract** |  |  |  |  |
| MIN7 (ARF-GEF) | AT3G43300 | 171 | 229 | 246 |
| HLB1 (TPR protein) | AT5G41950 | 32 | 27 | 45 |
| MIN13 (ARF-GEF) | AT3G60860 | 27 | 26 | 42 |
| ARFA1c | AT2G47170 | 7 | 6 | 6 |
| 14-3-3 CHI | AT4G09000 | 2 | 4 | 4 |
| Hsp81.4 | AT5G56000 | 3 | 3 | 2 |
| **Pulled down from detergent solubilized TGN vesicle protein preparation** |  |  |  |  |
| MIN7 (ARF-GEF) | AT3G43300 | 365 | 747 |  |
| MIN13 (ARF-GEF) | AT3G60860 | 115 | 205 |  |
| HLB1 (TPR protein) | AT5G41950 | 61 | 84 |  |
| SEC7-like ARF-GEF | AT4G38200 | 39 | 105 |  |
| SEC7-like ARF-GEF | AT1G01960 | 48 | 131 |  |
| MIN10 (14-3-3 KAPPA) | AT5G65430 | 2 | 3 |  |

<sup>a</sup>MIN7-GFP pull down was conducted using 1μM flg22-treated *min7/ MIN7-GFP* and Col-0 plants (as a negative control). Proteins detected with two or more peptides in both (for pull down with solubilized TGN vesicle protein preparation) or all three (for pull down with total leaf extract) replicates in *min7/ MIN7-GFP* plants, but not from Col-0 plants or Col-0/35S:*GFP* plants, are listed. Detected peptide numbers by mass spectrometry analysis are shown for each protein. Genes or proteins indicated by blue text are further analyzed in this study. Chloroplastic proteins LHCB4.1 [AT5G01530], FTSZ2-2 [AT3G52750], and pyridine nucleotide-disulphide oxidoreductase family protein [AT1G74470] were also detected in these pull downs, but were excluded for further analysis because MIN7 is normally localized in the TGN/EE. Proteins/genes analyzed in this study are shown in blue text.

We had previously developed specific antibodies against MIN7 and MIN10 [20], and therefore used these antibodies to perform targeted co-immunoprecipitation (co-IP) experiments with MIN7 and MIN10. As shown in Fig. S3, the physical interaction between MIN7 and MIN10 could also be detected in this assay. Thus, *in vivo* pull down and/or co-IP analyses showed that several host targets of HopM1 are components of a larger protein complex *in vivo*.

To further confirm protein-protein interactions among MIN proteins and to define interaction regions, we conduct yeast two-hybrid (Y2H) assays. As shown in Fig. 1C, the central and C-terminal regions of MIN7 (i.e., MIN7-M and MIN7-C) interacted with MIN10, and HLB1, respectively. Interestingly, however, unlike all MIN proteins [22], no interaction between HLB1 and tested fragments of HopM1 could be detected in Y2H assay (Fig. 1D), explaining why HLB1 was not identified in our initial Y2H screen [20] and suggesting that HLB represents a component of the MIN7 complex that is not targeted by HopM1.

### Genetic Analysis of the Components of the MIN7 Complex

As mentioned above, the *min7* mutant was previously shown to partially rescue the virulence defect of the ΔCEL mutant bacteria [20]. We investigated the possibility that mutants defective in genes encoding other components of the MIN7 complex might have a similar phenotype. For this purpose, we obtained T-DNA knockout mutant plants for *HLB1*, *MIN10*, *MIN13*, *14-3-3 CHI* and *ARFA1c* from the Arabidopsis Biological Resource Center and, after confirming their homozygosity by polymerase chain reaction (PCR) assays and lack of the corresponding transcripts by reverse transcriptase-PCR assays, subjected them to disease assays. The *hlb1* mutant (SALK_046760, *hlb1-5*, Figs. 2C-F) phenocopied the *min7* mutant and significantly rescued the virulence defect of the ΔEM mutant (Fig 2A). In contrast, *min10*, *min13, 14-3-3 chi* and *arfA1c* mutants were not more susceptible to the ΔEM mutant, compared with the wild-type Col-0 plants. The *hlb1* mutant also had a slightly increased susceptibility to the nonpathogenic *hrcC* mutant (Fig. 2B, which is defective in type III secretion [51], although the increased susceptibility to this strains was not as obvious as that to the ΔEM mutant bacteria. In contrast, the susceptibility of *hlb1* mutant plants to *Pst* DC3000 was similar to that of wild-type Col-0 plants.

**Figure 2.**
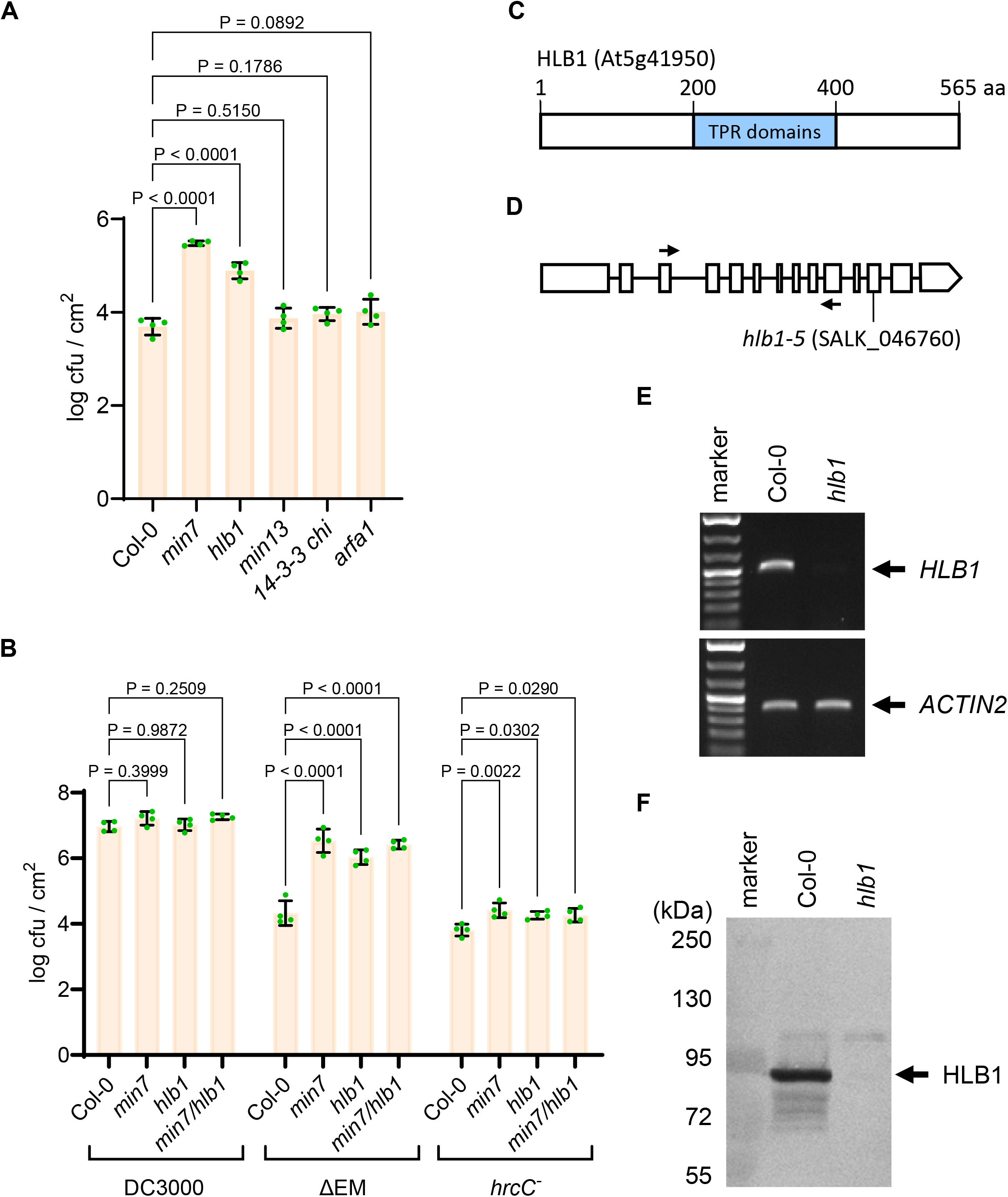
Analyses of Arabidopsis mutant plants defective in the components of the MIN7 complex. (A) Multiplication of the ΔEM mutant in *min7*, *hlb1*, *min13*, *14-3-3 chi*, and *arfA1c* mutant plants. T-DNA knockout lines of *min7*, *hllb1* mutant (SALK_046760), *min13* (SALK_033446), *14-3-3 chi* (SALK_142285) and *arfA1c* (SALK_136703) were dip-inoculated with the ΔEM mutant at 1×10^8^ cfu/ml and immediately covered with a clear plastic dome to maintain high humidity. Bacterial populations (mean ± SEM; n=4 leaf samples) in leaves were determined at day 4 post dip-inoculation. (B) Multiplication of *Pst* DC3000, ΔEM mutant and the *hrcC* mutant in *min7*, *hlb1*, *min7/hlb1* or Col-0 plants. Plants were inoculated by dipping with 1×10^8^ cfu/ml bacteria and immediately covered with a clear plastic dome to maintain high humidity. Bacterial populations (mean ± SEM; n=4 leaf samples) in leaves were determined at day 4 post dip-inoculation. Disease symptoms (chlorosis and necrosis) in Col-0 and mutant plants at day 4 are shown in Figure S4. (C) HLB1 contains multiple tetratricopeptide repeats. (D) The *hlb1-5* mutant (SALK_046760) carried a T-DNA insertion (vertical arrow) in the 12th exon. Two horizontal arrows indicate the positions of primers for RT-PCR analysis in E. (E) RT-PCR analysis of *HLB1* gene expression. No *HLB1* transcript was detected in the *hlb1* mutant after 25 cycles of RT-PCR. Col-0 plants were used as a positive control. (F) Western blot analysis of HLB1 in Col-0 and *hlb1* mutant plants using a HLB1-specific antibody developed in this study. The endogenous HLB1 protein was detected in Col-0, but not in the *hlb1* mutant.

To investigate whether there are additive effects between *min7* and *hlb1* mutations in disease phenotypes, we crossed the *min7* mutant to the *hlb1* mutant. The homozygous *min7/hlb1* double mutants were obtained. Disease assays showed that the *hlb1* mutation did not further increase the susceptibility of the *min7* mutant to the ΔEM mutant bacteria (Fig. 2B), providing genetic evidence that HLB1 and MIN7 likely act in the same pathway/complex. We also crossed the *min7* mutant to the *min10, min13* and *14-3-3 chi* mutants and obtained homozygous *min7/min10, min7/min13* and *min7/14-3-3 chi* double mutants. The susceptibility of these double mutants to the ΔEM mutant was similar to the *min7* single mutant (Fig. S5), suggesting that there is no further MIN7-dependent synergistic disease phenotypes for the *min10*, *min13* and *14-3-3 chi* mutants.

### HLB1 is localized to the MIN7-positive subcellular compartment in a MIN7-dependent manner

MIN7 and HLB1 were previously localized to the TGN/EE compartment in leaf and root cells, respectively [22, 23]. We were interested in determining whether MIN7 and HLB1 are co-localized in leaf cells. For this purpose, we transformed the *hlb1* mutant with the *35S::RFP-HLB1* fusion construct (Fig. 3A). As shown in Fig. 3B, *35S::RFP-HLB1* lines #2 and #3, which expressed the RFP-HLB1 fusion close to the endogenous HLB1 level, could complement the *hlb1* mutant and partially restored Arabidopsis resistance to the ΔEM mutant bacteria. This result suggests that the RFP-HLB1 fusion protein is functional. Surprisingly, confocal microscopic examination of transgenic Arabidopsis leaf cells showed RFP-HLB1 is localized diffusely in the cell in mock (H_2_O) treated leaves (Fig. 3C). Interestingly, however, we found that RFP-HLB1 becomes localized to both vesicle-like punctate structures and cytoplasm of leaf cells after flg22 treatment to activate pattern-triggered immunity (Fig. 3D).

**Figure 3.**
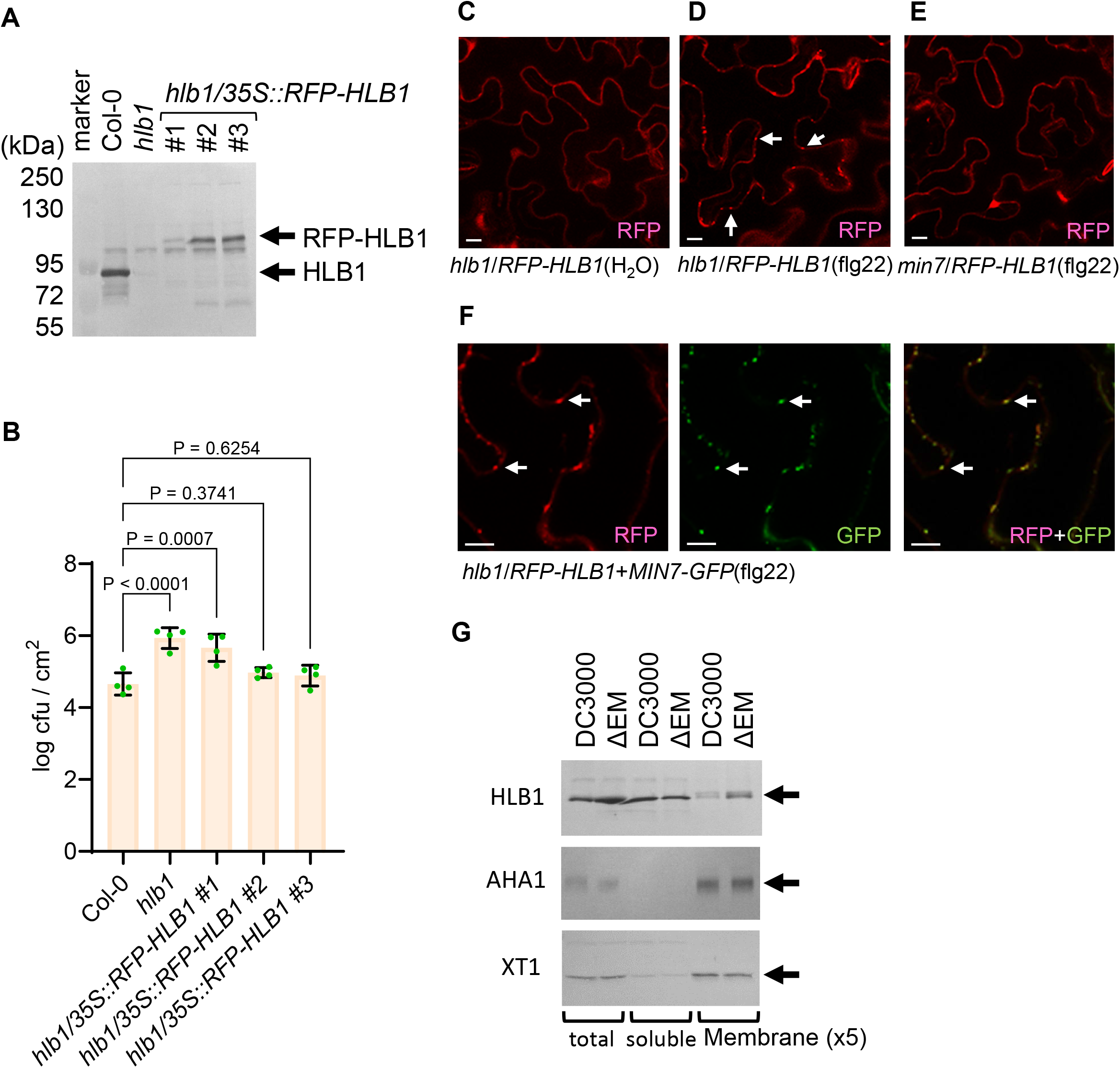
Localization of the HLB1 protein in Arabidopsis. (A) Western blot analysis of wild-type Col-0 and transgenic *hlb1* plants transformed with *35S::RFP-HLB1*. Total leaf extracts were separated by SDS-PAGE. HLB1 was detected by a HLB1-specific antibody. (B) Multiplication of the ΔEM mutant in Col-0, *hlb1* and *hlb1*/*35S::RFP-HLB1* plants. Arabidopsis plants were dip-inoculated with the ΔEM mutant at 1×10^8^ cfu/ml and immediately covered with a clear plastic dome to keep high humidity. Bacterial populations (mean ± SEM; n=4 leaf samples) in leaves were determined at day 4 post dip inoculation. (C) to (F) Representative confocal images of *hlb1/35S::RFP-HLB1* plants after H_2_O (C) or flg22 treatment (D), of *min7/35S::RFP-HLB1* plants after flg22 treatment (E), and of *min7*/*35S::RFP-HLB1*+*35S::MIN7-GFP* plants after flg22 treatment (F). 0.5 µM flg22 or H_2_O was hand-infiltrated into leaves 7 h before confocal microscope imaging. White scale bars: 10 μm. (G) HLB1 protein level during bacterial infection. Arabidopsis Col-0 leaves were hand-infiltrated with *Pst* DC3000 or the ΔEM mutant at 1×10^8^ cfu/ml and air-dried to let infiltrated leaves return to pre-infiltration appearance (∼1 h) and then covered with a clear plastic dome. Total proteins (T) were extracted from leaves 9 h after infection and separated into soluble (S) and total membrane (M) fractions by ultra-centrifugation. Protein samples were loaded onto a SDS-PAGE gel. HLB1, XT1, and AHA1 were detected with specific antibodies.

To determine whether the HLB1-localized vesicle-like structures MIN7-positive TGN/EE, we transformed *35S::RFP-HLB1* into *min7* plants expressing *35S:MIN7-GFP* [52] and examined subcellular localization of HLB1 and MIN7. As shown in Fig. 3F, HLB1-associated vesicle-like structures are MIN7-positve, suggesting that HLB1 and MIN7 are co-localized in the TGN/EE compartment. To further determine whether HLB1 localization to TGN/EE is dependent on MIN7, we transformed the *35S::RFP-HLB1* construct into the *min7* mutant. As shown in Fig. 3E, RFP-HLB1 signal was no longer found in vesicles even under flg22 treatment, but was completely diffused in the cytoplasm. This result suggests that, in the leaf cell, HLB1 is recruited to the TGN/EE compartment from the cytosol in a MIN7-dependent manner.

### Membrane-associated HLB1 is removed during *Pst* DC3000 infection in a HopM1-dependent manner

*Pst* DC3000-delivered HopM1 was shown to be sufficient to mediate the degradation of MIN2, MIN7 and MIN10 during infection [20]. In this study, we investigated whether HLB1, being a component of the MIN complex, is also degraded in a HopM1-dependent manner during *Pst* DC3000 infection. We found that the amount of membrane-associated HLB1 was reduced in *Pst* DC3000-infected leaves compared to that in ΔEM mutant-infected leaves, whereas the levels of HLB1 in the soluble faction were similar in DC3000- and ΔEM mutant-infected leaves (Fig. 3G). As controls, general membrane marker proteins such as the plasma-membrane protein AHA1 or the Golgi-associated protein XT1 was not reduced in the membrane fraction and their amounts were similar in DC3000- and ΔCEL mutant-infected leaves (Fig. 3G). This result suggests that it is the membrane-associated pool of HLB1 that is subjected to removal in a HopM1-dependent manner and is consistent with the HopM1 localization in the membrane organelle TGN/EE [22] and with the fact that HopM1 and HLB1 do not interact with each other in yeast cells (Fig. 1D). Therefore, it is possible that the reduction of membrane-associated pool of HLB1 during *Pst* DC3000 infection is likely caused by HLB1 dissociation from the TGN/EE upon MIN7 degradation by HopM1.

### *min7* and *hlb1* mutants have reduced immunity and increased water content in the apoplast

We previously showed that HopM1, when delivered from *Pst* DC3000, suppresses defense-associated callose deposition [12], and that *min7* plants are compromised in pathogen-associated molecular pattern (PAMP)-, effector-, and BTH-triggered immunity [22]. In this study, we investigated the possibility that the *hlb1* mutant may have similar immune phenotypes. Indeed, as shown in Fig. 4A, bacterial populations in the *hlb1* mutant were significantly higher compared to Col-0 plants during PAMP (flg22)- and BTH-triggered immunity. And, in *hlb1*, callose deposition under PTI-immunity was significantly lower than that in Col-0 (Fig. 4B).

**Figure 4.**
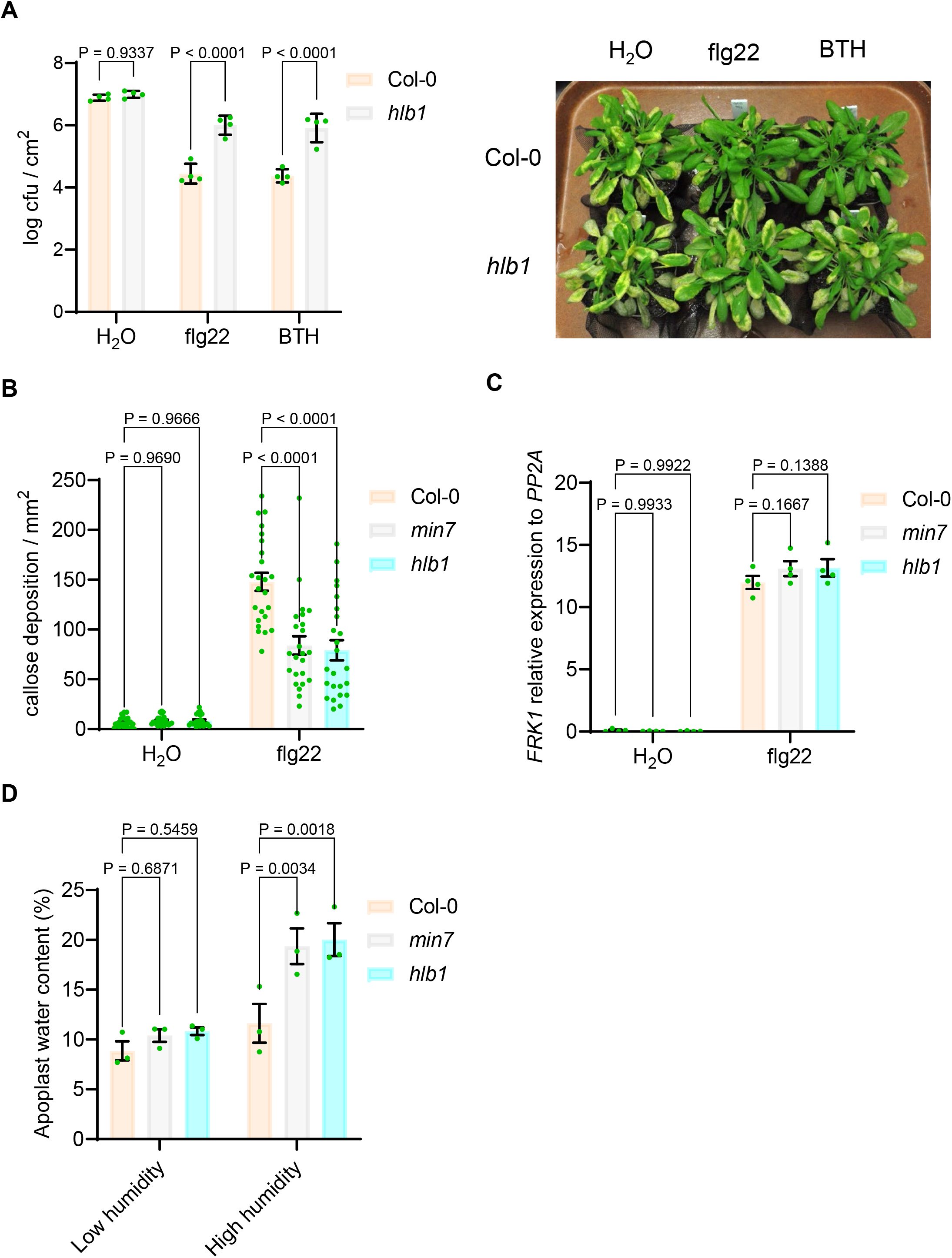
Role of HLB1 in effector-, BTH- and flg22-triggered immunity. (A) Multiplication of *Pst* DC3000 in Col-0 and *hlb1* plants which pre-treated with flg22 or BTH. H_2_O, 1 µM flg22 or 30 µM BTH was spray to plants and covered by a clear plastic dome for 24 h before dip inoculation with *Pst* DC3000 at 1×10^8^ cfu/ml. After dip inoculation, plants were covered by a clear plastic dome to keep high humidity. Bacteria populations (mean ± SEM; n=4 leaf samples) in leaves were determined at day 3 post dip inoculation. Disease symptoms (chlorosis and necrosis) in Col-0 and *hlb1* plants were recorded at day 4. (B) and (C) Callose deposition and *FRK1* gene expression in *hlb1* leaves. Arabidopsis leaves were syringe-infiltrated with 1 µM flg22 and air-dried to let infiltrated leaves returned to pre-infiltration appearance (∼1 h) and then covered by a clear plastic dome. Callose deposition (B) was observed at 8 hr post flg22 inoculation (mean ± SEM; n=24 callose stained leaf pictures in mm^2^). The *FRK1* gene expression (C) was detected at 8hr hr post flg22 inoculation (mean ± SEM; n=4 leaf samples). (D) Apoplastic water content in *hlb1* mutant leaves. Five week-old Arabidopsis plants were moved into control (∼50%) or high (∼99%) humidity setting chamber. Apoplastic water contents were determined at 24hr after humidity change (mean ± SEM; n=3 biological repeats).

We noted that the host immune responses affected by HopM1 (delivered by *Pst* DC3000 during infection) appear to be limited to callose deposition [12, 20, 22]. Gene expression analyses by Northern blot or semi-quantitative reverse transcription-polymerase chain reaction (RT-PCR) assays did not reveal an effect of *Pst* DC3000-delivered HopM1 or the *min7* mutation (which partially mimics the virulence activity of HopM1) on canonical salicylic acid- and flg22-triggered defense marker gene expression [12]. In this study, we confirmed lack of an effect of the *min7* and *hlb1* mutations on defense the *FRK1* marker gene expression elicited by flg22 using quantitative PCR assay, which is considered to be more quantitative than Northern blot and RT-PCR assays (Fig. 4C). These results suggest that the previously observed effect of HopM1 and the *min7* mutation on callose deposition and bacterial multiplication during PAMP (flg22)-triggered immunity, is unlikely through their direct effects on canonical defense gene expression.

While it remained unclear why bacteria-delivered HopM1 and the *min7* mutation affects callose deposition and immunity without affecting defense gene expression, it was recently discovered that, besides suppressing defense-associated callose deposition in the apoplast, HopM1 and AvrE share another virulence activity: induction of an aqueous apoplast (water-soaking) under high humidity [16-18]. This new result raised the possibility that the callose suppression phenotype associated with HopM1 and the *min7* mutant may be related to altered water homeostasis in the apoplast. To test this hypothesis, we directly measured the apoplast water content in the *min7* and *hlb1* mutant leaves. We found that the *min7* and *hlb1* mutant leaves contain twice as much water as the wild-type Col-0 plants under high humidity condition (Fig. 4D).

Strikingly, artificial water supplementation to the apoplast alone was sufficient to compromise callose deposition under PAMP (flg22)-triggered immunity [53], suggesting that higher water content in *min7* and *hlb1* is likely responsible for reduced callose deposition in response to flg22 elicitation of pattern-triggered immunity.

## Discussion

In the past decade, efforts to identify the host targets of pathogen effectors have taken center stage in the study of microbial pathogenesis in plants [3, 5-8]. An important finding from these studies is that, at least for the model pathogen *Pst* DC3000, many type III effectors appear to target various components of the plant immune system. However, in many cases multiple host targets have been reported for a single effector [1]. It is often not clear whether all of them are biologically relevant to the virulence functions of the cognate effectors during pathogen infection, and, if they are, whether they are involved in the same or distinct host cell functions. HopM1 is a good example of a single effector targeting multiple, sequence-unrelated host proteins [18, 20, 27]. In this study, we found that MIN7 forms complex with several other MIN proteins and identified other candidate components of the MIN7 complex. Remarkably, HopM1 could mediate the degradation of several components of the MIN7 complex, including MIN2, MIN7, and MIN10, as shown previously [20], and reduce the membrane-associated pool of the newly identified HLB1 protein (this study). These results provide a satisfactory explanation for the ability of HopM1 to target multiple, sequence-unrelated host targets and, more importantly, uncovered an infection-relevant ARF-GEF-based protein complex in plants (Fig. 5).

**Figure 5.**
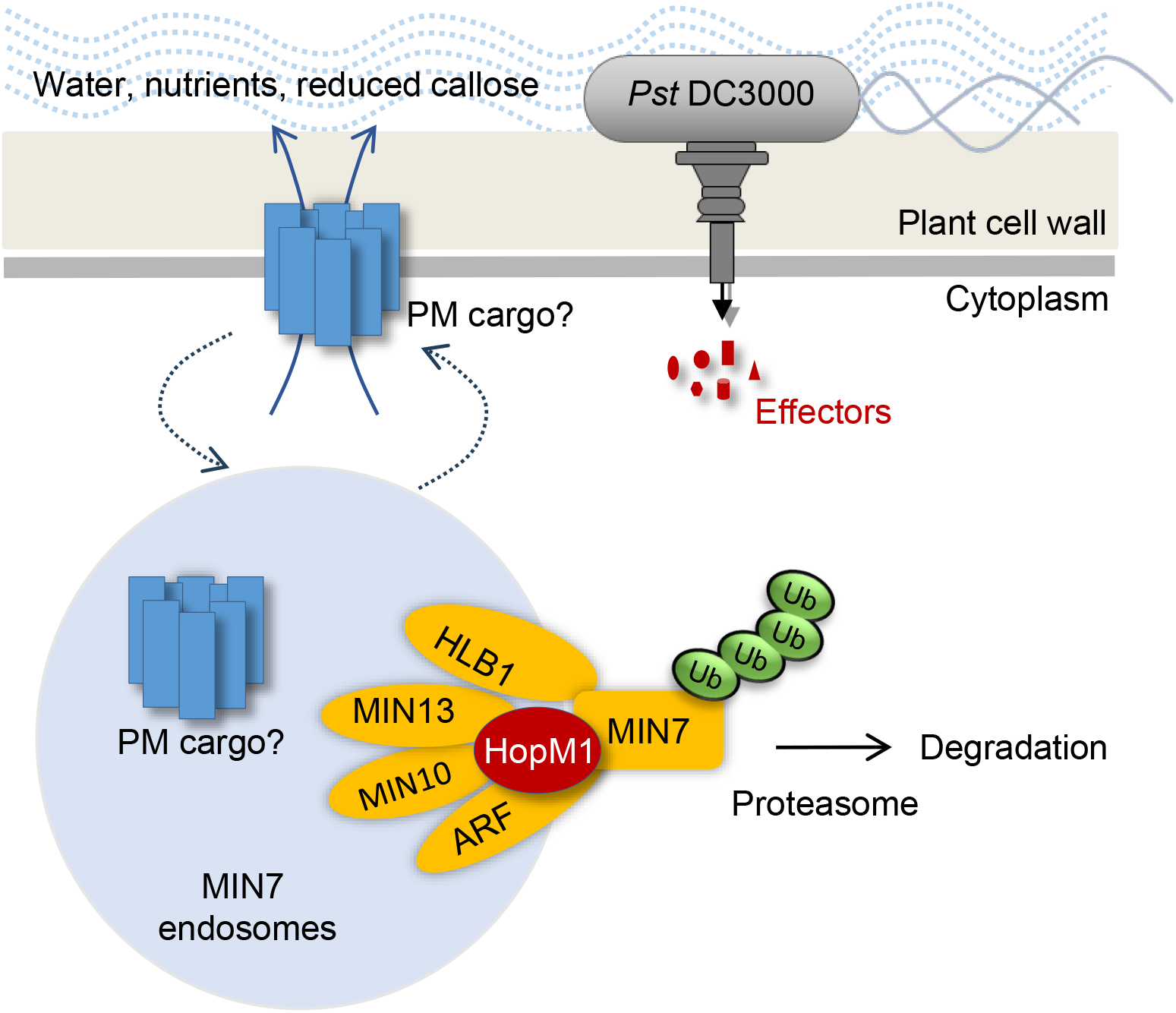
A diagram of the MIN7 complex in the TGN/EE involved in plasma membrane (PM) cargo recycling and apoplastic water content and immunity. Components of the MIN7 complex are shown. The MIN7 complex is involved in controlling apoplastic water content and defense activity.

In this study, we provide evidence that, like the *min7* mutant, the *hlb1* mutant could partially restore the infectivity of the ΔCEL mutant, which lacks the *hopM1* gene. Also like the *min7* mutant, the *hlb1* mutant could not fully restrict bacterial multiplication during PAMP-, effector- and BTH-triggered immunity. In contrast, *MIN10* (i.e., *14-3-3 KAPPA)*, *MIN13* (i.e., *ARF-GEF, BIG2)*, *14-3-3-CHI,* and *ARFA1c*, knockout mutants did not show such phenotypes. We noted that HLB1 is a unique protein, whereas the others belong to multi-gene families (Fig. S2). Therefore, a lack of disease phenotype in *min10*, *min13*, *14-3-3 chi* and *arfA1c* mutants could be due to functional redundancy or compensatory effects among members of these particular gene family members. Indeed, 14-3-3 KAPPA and RAD23 (MIN2) proteins have recently been shown to be the host targets of other effectors from *Pst* DC3000 and other pathogens [54-58], and a study showed that chemical inhibitors of client protein interaction with the 14-3-3 family of proteins could rescue the virulence defect of the *Pst* DC3000 ΔCEL mutant [25]. This observation suggests that, as a group, 14-3-3 proteins may be biologically relevant targets of HopM1. Thus, it is likely that multiple components of the MIN7 ARF-GEF complex are involved in diverse plant-bacterial interactions.

Previous studies showed that HopM1 and/or MIN7 are localized in the TGN/EE [22, 23]. Although we have not determined the subcellular localization of all MIN7-interacting proteins, the observation that they interact with MIN7 indicates that at least a pool of each of these MIN7-interacting proteins is associated with the TGN/EE. We previously showed that the MIN7 protein level is significantly enhanced during immune response [22]. Here we show that that the increased MIN7 protein level leads to the recruitment of HLB1 to the TGN/EE (Fig. 3). Taken together, our results suggest that the assembly of the MIN7 complex at the TGN/EE may be responsive to immune stimulation, at least in the leaf tissue. Interestingly, only membrane-associated HLB1, but not soluble HLB1, is reduced in a HopM1-dependent manner during *Pst* DC3000 infection. The HopM1-depedenent reduction of the membrane-associated pool of the HLB1 protein is likely indirect because, unlike MIN proteins, we could not detect an interaction between HopM1 and HLB1 in Y2H (Fig. 1) and because the localization of HLB1 to membrane/vesicles is dependent on MIN7 (Fig. 3). HopM1-mediated degradation of MIN7 in the TGN/EE therefore could cause TGN/EE-localized HLB redistributed to the cytosol.

HopM1 is one of the most conserved effectors in *P. syringae* [9], but shares no sequence similarity to any protein of known function. How HopM1 mediates degradation of proteins of the MIN7 complex will require future study. We find it interesting that one of HopM1’s host targets, MIN2 (RAD23a, which shuffles ubiquinated proteins to the proteasome) was not detected as a component of the MIN7 complex in this study. It is possible that, different from other MIN proteins, MIN2 may not be a component of the MIN7 complex, but is recruited by HopM1 as part of a novel mechanism to induce proteasomal degradation of proteins of the MIN7 complex (Fig. 5). Consistent with this possibility, although MIN7 does not appear to interact with any components of the ubiquitination/proteasome system in our previous Y2H screen and the *in planta* pulldown screen in this study, HopM1 could pull down a number of components of the ubiquitination/proteasome system, in addition to MIN7 and MIN10 proteins, when transiently overexpressed in *Nicotiana benthamiana* leaves [26]. Thus, HopM1 could be a novel adaptor that connects the MIN7 protein complex to the host ubiquitination/proteasome system.

Previous studies implicated HopM1 in modulating two apparently distinct host cellular processes: immune modulation [19] and apoplastic water-soaking [16]. Curiously, however, the host immune responses affected by HopM1 (when delivered by *Pst* DC3000 during infection) appear to be limited to apoplastic callose deposition [12, 20, 22], whereas other immune-suppressing bacterial effectors often affect multiple defense responses, including defense gene expression [59-61]. Furthermore, in an experiment to define the minimal functional repertoire of *Pst* DC3000 effectors, Cunnac and colleagues (2011) showed that HopM1’s virulence function is different from that of canonical immune-suppressing effectors, such as AvrPto [62]. Our disease reconstitution experiments using Arabidopsis host target mutants supported this notion [16]. These studies led to a new model for *Pst* DC3000 pathogenesis in susceptible plants: *Pst* DC3000 translocates at least two functional groups of effectors into the host cell: one group, exemplified by AvrPto, suppresses immune signaling [61, 63], whereas the other group, such as HopM1, induces the formation of an aqueous apoplast [16, 64]. In this and a related study [53], we provide evidence that these two distinct pathogenic processes may influence each other at some level. In particular, we found that the *min7* and *hlb1* mutant leaves contain higher water content (this study) and transient apoplast water supplementation, which mimics HopM1-mediated water-soaking during infection, was sufficient to suppress callose deposition elicited by flg22 [53] thereby phenocopying characteristic effects of HopM1 on immune responses. There seems to be “cross-talk” between immune responses and water availability in the apoplast. We speculate that an increased water content in the *min7* and *hlb1* mutant leaves could not only facilitate the flow of water and nutrients to bacteria, promote the spread/egression of bacteria, but also may negatively affect the deployment of apoplastic some host defense responses. Collectively, these effects may explain the compromised ability of *min7* and *hlb1* mutants in restricting bacterial multiplication during PAMP-triggered immunity.

In summary, by following the virulence action of the bacterial effector HopM1, we have identified multiple components of a TGN/EE-localized ARF-GEF-based protein complex that plays an important role in mediating *Pst* DC3000 pathogenesis. Central to this complex appears to be the MIN7 ARF-GEF, which engages physical interactions with other components via distinct regions (Fig 1 and 5). Although HopM1 was the first bacterial effector shown to destroy an ARF-GEF, via the host proteasome [20], the type III effector EspG from the human pathogen *Escherichia coli* O157:H7 also targets host ARF-regulated trafficking [65]. In addition, the *Legionella pneumophila* effector RalF secreted through the type IV secretion system is itself an ARF-GEF [66]. Importantly, MIN7 has recently been shown to be required for Arabidopsis and wheat resistance against the fungal pathogen *Fusarium graminearum* [67]. These findings suggest an emerging paradigm of ARF- and ARF-GEF-based battles between eukaryotic hosts of different kingdoms and pathogens. As such, further characterization of the MIN7 complex and how HopM1 attacks this complex should provide insights into a fundamental, yet poorly understood aspect of host-pathogen battles.

## Materials and Methods

All experiments reported in this paper were performed two or more times with similar results.

### Bacterial strains, plasmids, and media

*P. syringae* strains used are wild-type *Pst* DC3000 [68], the ΔCEL mutant [10], the ΔEM mutant [21], the *hrcC* mutant [51]. Bacteria were grown in low-salt Luria-Bertani medium at 28°C with antibiotics. Antibiotics were used at the following concentrations (µg/ml): ampicillin, 200; rifampicin, 100; spectinomycin, 50; tetracycline, 30; kanamycin 50.

### Plant growth and bacteria enumeration

Methods for plant growth and bacteria inoculation and enumeration were described previously [22]. Five-week-old plants were used for experiments. Infected plants were monitored daily over a 3- to 4-day period for symptom development and bacterial multiplication.

### Construction of fusion genes in binary vector

The *HLB1* coding sequence was amplified by PCR using the following primers. The amplified *HLB1* fragment was cloned into pENTR/D cloning vector (Invitrogen), and transferred by LR recombination into the binary expression vector pGWB555 [69] to generate a *35S::RFP-HLB1* fusion.

*HLB1*:

Sense primer: <u>ATG</u>GCGGATACTGTTGAAGAG (start codon underlined)

Antisense primer: <u>TTA</u>ACCGGTGATAATACCGGC (stop codon underlined)

### Production of transgenic Arabidopsis

The binary vectors containing *35S::RFP-HLB1* fusion gene was introduced into *Agrobacterium tumefaciens* C58C1 by electroporation. Arabidopsis plants were transformed using the floral dip method [70]. Hygromycin was used to select for transgenic T1 plants, which were further screened by western blot using HLB1 specific antibody. Homozygous T3 plants expressing fusion were used for analyses.

### Confocal microscopy analysis and imaging

Leaf pieces of stable transgenic Arabidopsis plants were mounted in water. Imaging was conducted with an Olympus FluoView FV1000 Laser Scanning Confocal Microscope (Tokyo, Japan) configured on an inverted IX81 automated microscope with Argon ion laser for GFP at 488 nm. A Helium Neon laser was used to visualize red fluorescent proteins at 543 nm. All imaging experiments were performed with a 40X PlanApo N oil immersion objective (NA 1.42). Images were processed using Olympus Fluoview Viewer software and Adobe Photoshop Elements (Adobe).

### RT-PCR and RT-qPCR analysis for gene expression

Total RNA was purified from leaf tissues by RNeasy Mini Plant Kit (Qiagen). First-strand cDNAs were synthesized from 200 ng of total RNA by using oligo dT primer and M-MLV Reverse Transcriptase (Invitrogen) according to the manufacturer’s instructions. PCR amplification was carried out using the following primers for 25 cycles.

*HLB1*:

Sense primer: TGCATTATACAACTGGGCACT

Antisense primer: GCATGTCTTTGCCGTTGCCTG

*ACTIN2*:

Sense primer: TGTGTCTCACACTGTGCCAAT

Antisense primer: CTTCCTGATATCCACATCACA

qPCR was run on the 7500 Fast Real-Time PCR system (Applied Biosystems) with SYBR master mix (Applied Biosystems) and following primers.

*FRK1*:

Sense primer: CTTCCATCGAGGTACAAAGATGAC

Antisense primer: CAGTGCTCATGACAGTAGAAGC

*PP2A*:

Sense primer: GGTTACAAGACAAGGTTCACTC

Antisense primer: CATTCAGGACCAAACTCTTCAG

### Preparation of the HLB1 specific antibody

The *HLB1* coding sequence corresponding to the N-terminal 200 amino acids was amplified by PCR using the following primers and was cloned into pGEM-T-easy vector (Promega). The *Nde*I and *Xho*I fragment of *HLB1_1-200_* was cloned into pET28a vector (Novagen) and was transformed into *E*. *coli* BL21(DE3). The 6×His::HLB1_1-200_ protein production was induced by adding 0.5 mM IPTG to LB bacterial culture for 3 h at 37 L. HLB1 protein was purified from total cell lysate by using Ni-NTA agarose beads in the extraction buffer (50 mM Tris-Cl, pH 8.0, 250 mM NaCl, 5 % Glycerol, 0.1 mM PMSF). Purified 6×His::HLB1 protein was injected to rabbit to raise HLB1 specific antibody (Cocalico Biologicals, Inc).

Sense primer: <u>CATATG</u>GCGGATACTGTTGAAGAG (*Nde*I site underlined, start codon double underlined)

Antisense primer: <u>CTCGAGTTA</u>AGATCGACCCTCTTCGTCGTTAAC (*Xho*I site underlined, stop codon double underlined)

### HLB1 detection under bacterial infection

Arabidopsis Col-0 leaves were infiltrated with 1×10^8^ cfu/ml of *Pst* DC3000 or ΔEM mutant. Ten hours later, treated leaves were harvested and homogenized in homogenization buffer (50mM HEPES-KOH, pH 7.5, 250mM NaCl, 0.5mM EDTA, 1mM PMSF). The homogenate was centrifuged at 4°C for 15 min at 10,000×*g* and the supernatant was filtered through Miracloth (Calbiochem) to remove plant debris. The filtered supernatant (total extract) was centrifuged again at 100,000×*g* for 1 h at 4°C to yield soluble (supernatant) and membrane (pellet) protein fractions. Fractionated proteins were separated on NuPAGE 4-12% Bis-Tris mini gels (Life Technology) and transferred onto Immobilon-P membranes (Millipore) for immunoblotting analysis. The amount of membrane fraction sample was 5-fold higher than that of other two fractions. HLB1, PM-localized H^+^-ATPase (AHA1), and Golgi-localized xyloglucan xylosyltransferase (XT1) were detected with anti-HLB1, anti-AHA1 (abcam), and anti-XT1 (gift from Dr. Ken Keegstra), respectively.

### TGN vesicle preparation

TGN protein purification was followed previous report [71]. Four week-old Col-0 and *min7* plants were sprayed with 1µM flg22. Six hours later, Arabidopsis leaves were collected and gentry homogenized in TGN isolation buffer (50mM HEPES-KOH pH7.5, 0.45M Sucrose, 1mM EDTA, 0.5% PVP#10,000, 1mM DTT, 1% Protease Inhibitor Cocktail, 1% Phosphatase Inhibitor Cocktail, 100µM MG132, 30µM BFA). Homogenized plant extract was centrifuged at 10,000×*g* for 15 min at 4°C to remove insoluble materials. Supernatant was filtered through Miracloth (Calbiochem) to remove plant debris and was centrifuged again at 100,000×*g* for 1 h at 4°C. TGN vesicle extracts were collected in supernatant.

### Native PAGE analysis of MIN7 complex

Native PAGE analysis was done with using NativePAG NovexR Bis-Tris Gel System (Life Technologies). TGN vesicle crude extract was detergent-solubilized in Native PAGE Sample Buffer (50 mM BisTris, 50 mM NaCl, 10% w/v Glycerol, 0.001% Ponceau S, 1% DDM, pH 7.2) and applied on NativePAG NovexR 4–16% Bis-Tris Gel. After electrophoresis, separated proteins were transferred to Immobilon-P membrane (Millipore). Protein transferred membrane was dried and washed with methanol to remove extra Coomassie blue before western blot analysis. MIN7 complex was detected with MIN7 specific antibody [22].

### Pull-down and mass spectrometry analysis of the MIN7 complex

Four week-old *min7*/*35S::MIN7-GFP*, Col-0/*35S::GFP*, and Col-0 plants were sprayed with 1µM flg22. Six hours later, Arabidopsis leaves were collected and homogenized in detergent-containing binding buffer (10mM Tris-HCl pH7.4, 150mM NaCl, 0.5mM EDTA, 0.2% NP-40, 1% Protease Inhibitor Cocktail, 1% Phosphatase Inhibitor Cocktail). Total protein extracts were collected after centrifugation at 10,000×*g* for 15 min at 4°C to remove insoluble materials. Supernatant was filtered through Miracloth (Calbiochem) to remove plant debris and was incubated with GFP-trap A beads (ChromoTek) with gentle shaking for 2 hour at 4°C. Beads were washed three times with washing buffer (10mM Tris-HCl pH7.4, 150mM NaCl, 0.5mM EDTA). MIN7 complex proteins were digested on beads and analyzed by LC/MS/MS at the MSU Proteomics Core Facility. We also pull-down MIN7 complex from purified TGN vesicle extract. TGN vesicle extracts from flg22 treated *min7*/*35S::MIN7-GFP* and Col-0 plants were incubate with GFP-trap A beads for 3 hour at 4°C. Beads were washed three times with detergent-containing TGN washing buffer (50mM HEPES-KOH pH7.5, 0.25M Sucrose, 1mM EDTA, 150mM NaCl, 0.1% NP-40).

### Yeast two-hybrid (Y2H) analysis

The LexA-based Y2H system (Clontech) was used for protein interaction analysis. Plasmid constructs expressing the full-length HopM1 (aa1-712; BD::*hopM1*), the N-terminal region (aa1-300; BD::*hopM1-N*), the C-terminal region (aa301-712; BD::*hopM1-C*) were made. Full-length RAD23 (AD::*MIN2*) and full length 14-3-3 kappa (AD::*MIN10*) were made in previous studies [11, 20]. The full-length *HLB1* and truncated *MIN7* fragments (see below) were cloned into pENTR/D cloning vector (Invitrogen), and transferred by LR recombination into pB42AD-GW and pGILDA-GW (Clontech), respectively. The following primers were used:

AD::*HLB1*:

Sense primer: <u>ATG</u>GCGGATACTGTTGAAGAG (start codon underlined)

Antisense primer: <u>TTA</u>ACCGGTGATAATACCGGC (stop codon underlined)

BD::*MIN7-N* (aa1-570):

Sense primer: <u>ATG</u>GCGGCTGGTGGATTTTTGACTCGA (start codon underlined)

Antisense primer: <u>TTA</u>TGGGACATCTTCCCTGCTTTTGGTTTC (stop codon underlined)

BD::*MIN7-M* (aa571-1207, including the Sec7 catalytic domain):

Sense primer: AGCAACTTTGAGAAGGCTAAAGCTC

Antisense primer: <u>TTA</u>TCCCTCTGCAAGCCGATCCTCACAT (stop codon underlined)

BD::*MIN7-C* (aa1208-1739):

Sense primer: CTTATACCCGGTGGTGTTCTTAAGCC (start codon underlined)

Antisense primer: <u>TTA</u>CTGTTGCAAAAGTGGCTTCAATTG (stop codon underlined)

Plasmid constructs were transformed into the EGY48 strain carrying the *lacZ* reporter plasmid. For western blot analysis, yeast strains were grown in galactose-inducing media for 16 hr. Total yeast proteins were extracted in 2×SDS sample buffer, separated on NuPAGE 4-12% Bis-Tris mini gels, and transferred onto Immobilon-P membranes (Millipore) for immunoblotting. AD and BD fusion proteins were detected with a chicken HA epitope antibody (Aves Labs) and a rabbit LexA binding domain antibody (Clontech), respectively.

### Detection of apoplastic water content

Five week-old Col-0, *min7*, and *hlb1* plants were moved into a control (∼50%) or high (∼99%) humidity chamber and kept for 24 hr. Eight leaves were infiltrated with 50 µM indigo carmin solution. After removing extra solution on leaf surfaces, apoplastic fluid was corrected by centrifuged at 2,000 ×g for 5 min at 4 °C. Absorbance of indigo carmine at 610 nm in the original indigo carmine solution vs. in the recovered apoplastic fluid were determined by SpectraMax M2 (Molecular Devices). Apoplastic water content was calculated following a previous study [72].

## Author contributions

KN and SYH designed research; KN, LI performed research; HT contributed materials; KN analyzed data; KN and SYH wrote the manuscript.

## Acknowledgements

We thank members of the He laboratory for stimulating discussions and Kyaw Aung for critical reading. We are thankful to Dr. Melinda Frame of Center for Advanced Microscopy for assistance with confocal microscopy and to Doug Whitten of the MSU Proteomics Core Facility for assistance with Mass Spectrometry and data analysis.

## Funding

National Institutes of Health (NIAID 1R01AI155441 (S.Y.H.)

Duke Science and Technology Initiative (S.Y.H.)

## Supporting information

Supporting Table (Table S1)

Supporting Figures (Figs. S1 to S5)

## Supporting Information

**Table S1. Arabidopsis proteins pulled down with GFP *in vivo*.**

**Figure S1.**
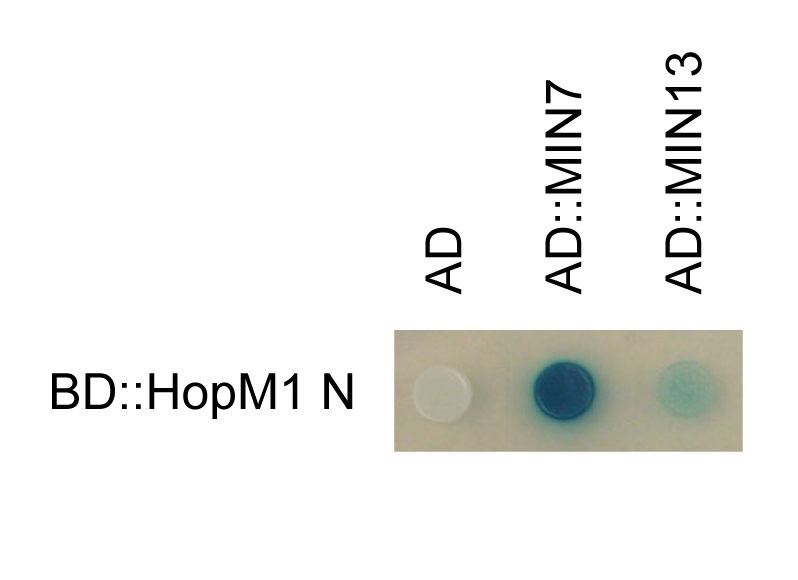
HopM1 weakly interacts with MIN13 (an ARF-GEF) in Y2H assay. HopM1-N and MIN13 were expressed from pGILDA (BD fusion) and pB42AD (AD fusion), respectively. Yeast cultures were spotted and grown on minimal medium containing galactose and X-gal. A blue color indicates interaction, whereas a white color indicates no interaction. AD-MIN7 was used as a positive control.

**Figure S2.**
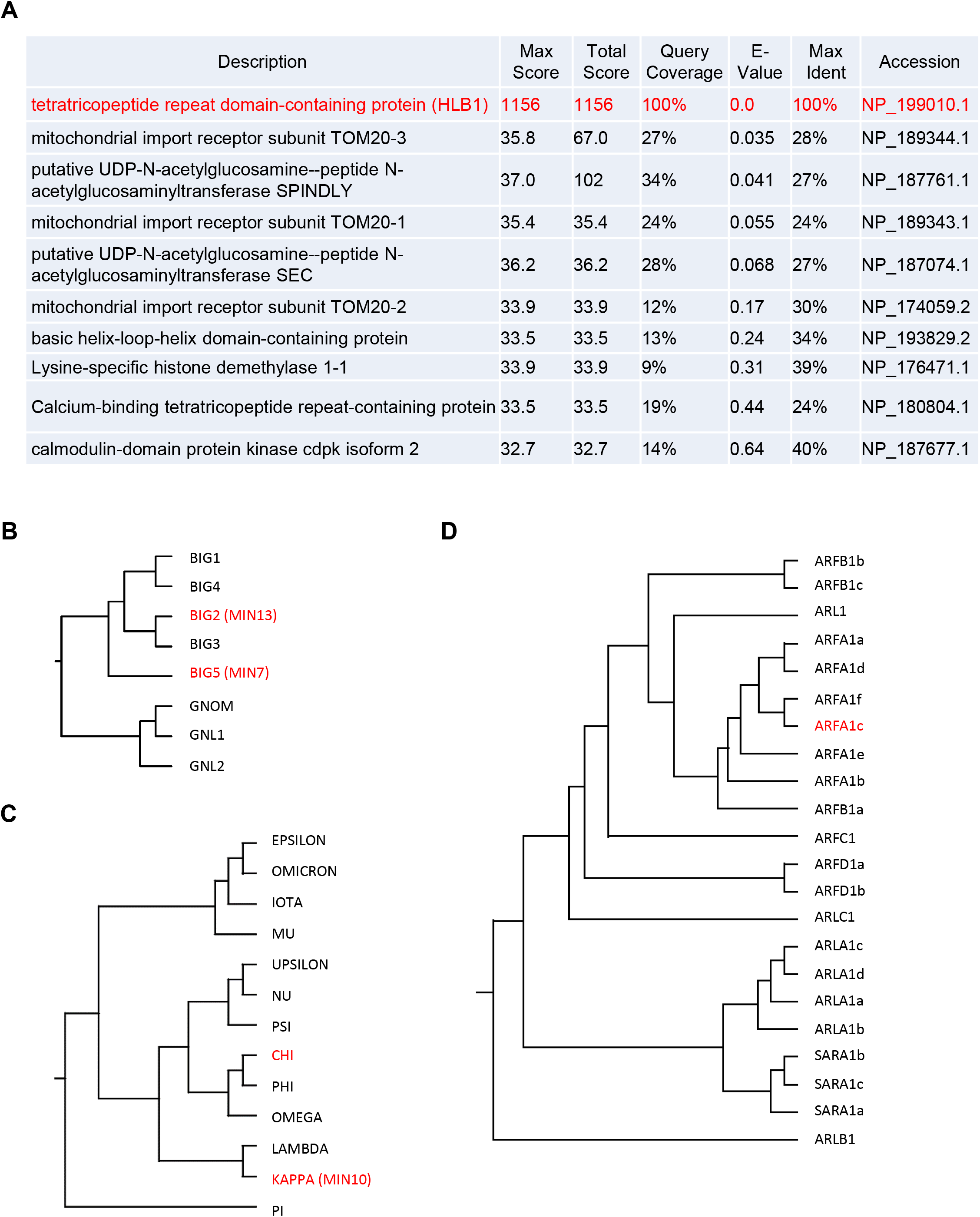
Gene families of the components of the MIN7 complex. (A) HLB1 does not share significant sequence similarity to any protein of known function. BlastP search against the nr database was performed using HLB1 as a query. The names and BlastP scores of the top ten protein sequences are displayed. (B) A phylogenetic tree of ARF-GEF proteins constructed using CLUSTAL 2.1 Multiple Sequence Alignments. (C) A phylogenetic tree of 14-3-3 proteins constructed using CLUSTAL 2.1 Multiple Sequence Alignments. (D) A phylogenetic tree of ARF GTPase proteins constructed using CLUSTAL 2.1 Multiple Sequence Alignments.

**Figure S3.**
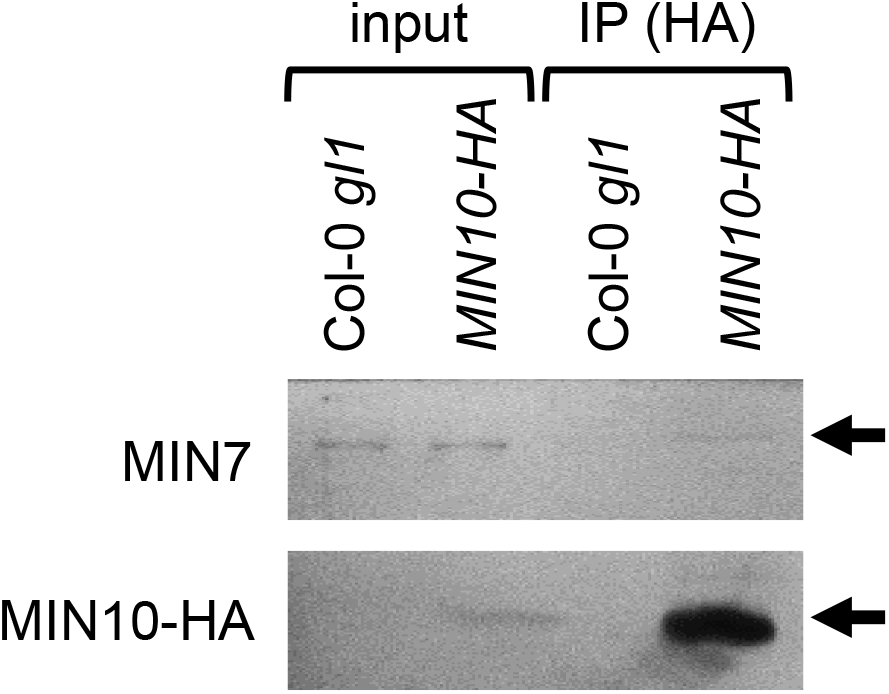
Co-IP analysis of MIN7 and MIN10 proteins. *35S::MIN10-HA*/Col-0 and Col-0 plants were treated with 1µM flg22 for 6 h. Total Arabidopsis extracts (Input) and immunoprecipitates (HA-agarose IP) were separated on an SDS-PAGE gel and subjected to immunoblot analysis. MIN7 and MIN10-HA were detected by MIN7- and HA-specific antibodies, respectively.

**Figure S4.**
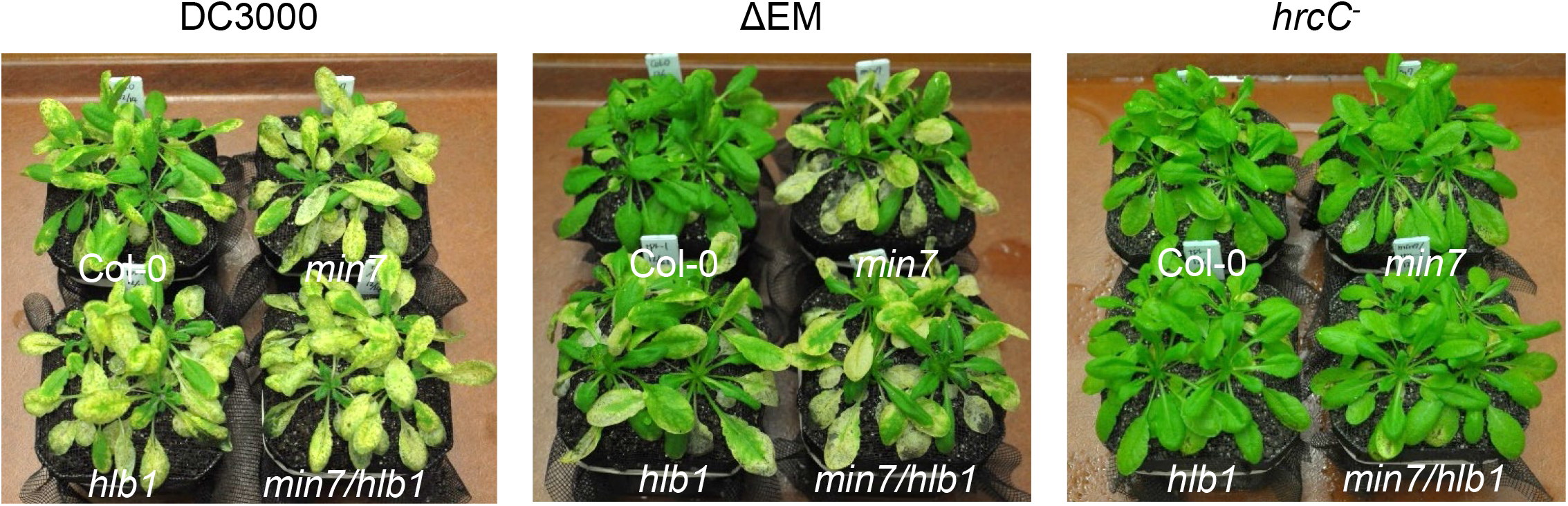
Disease symptoms on *min7*, *hlb1*, *min7/hlb1* or Col-0 plants. Plants were inoculated by dipping with 1×10^8^ cfu/ml bacteria and immediately covered with a clear plastic dome to maintain high humidity. Disease symptoms (chlorosis and necrosis) in Col-0 and mutant plants were recorded at day 4.

**Figure 5.**
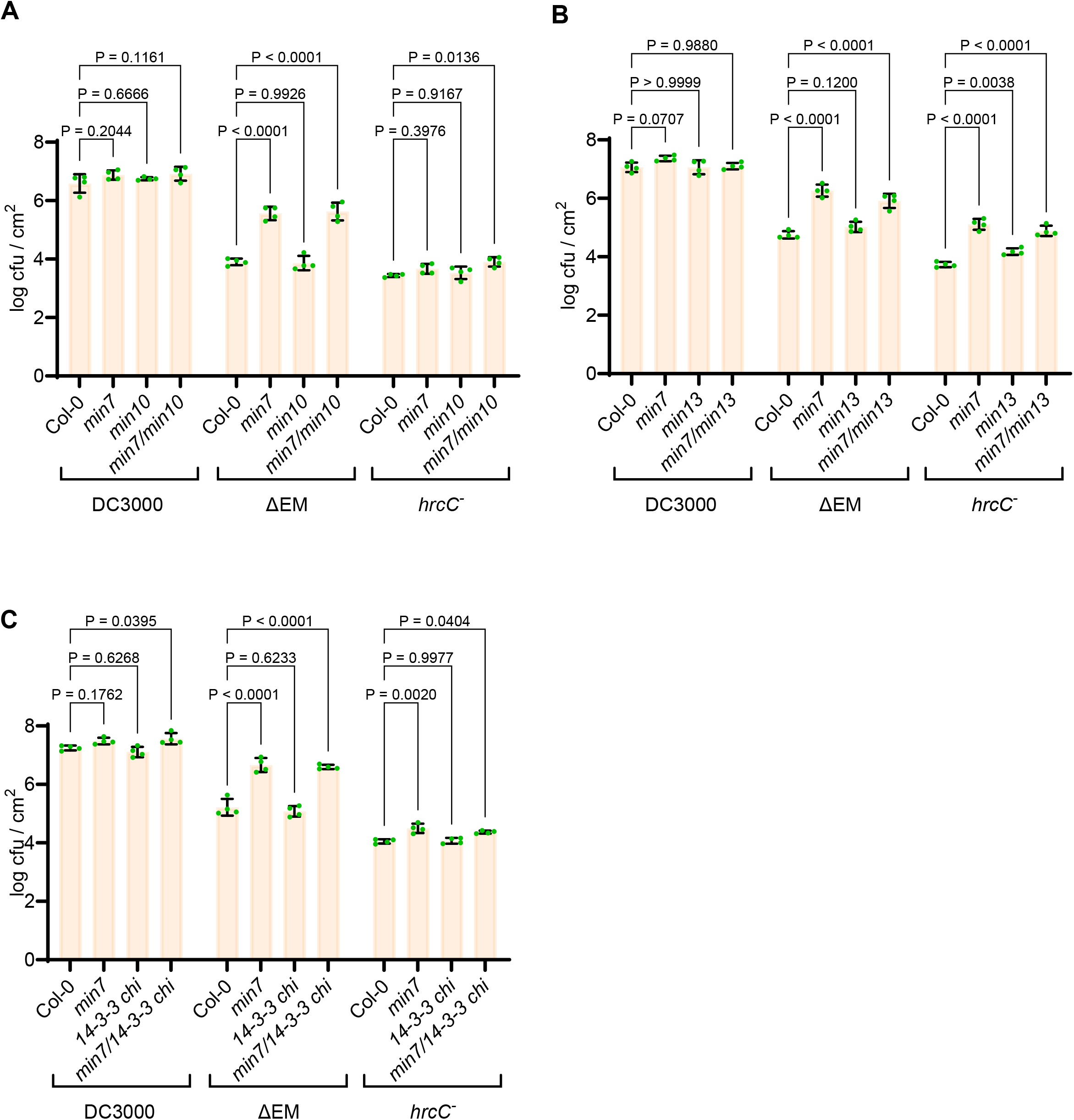
Bacterial multiplication in Arabidopsis mutant plants affected in the components of the MIN7 complex. (A) to (C) Arabidopsis plants were dip-inoculated with the *Pst* DC3000, ΔEM mutant, and the *hrcC* mutant at1×10^8^ cfu/ml and immediately covered with a clear plastic dome to maintain high humidity. Bacterial populations (mean ± SEM; n=4 leaf samples) in leaves were determined at day 4 post dip inoculation.

