## Supplemental Table 1 for "Multiple host targets of *Pseudomonas* effector protein HopM1 form a protein complex regulating apoplastic immunity and water homeostasis"

**Table S1. Arabidopsis proteins pulled down with GFP *in vivo^a^*.**

| **Identified Proteins** | **Accession #** | **#1** | **#2** | **#3** |
| --- | --- | --- | --- | --- |
| **Pulled down from total leaf extract**  GFP | AT3G43300 | 52 | 71 | 48 |
| Di-haem cytochrome | AT2G07727 | 1 | 1 | 1 |

*^a^*GFP pull down was conducted using 1µM flg22-treated Col-0/*35S::GFP* plants and Col-0 plants (as a negative control). Proteins detected in all three pull down replicates in Col-0/*35S::GFP* plants, but not from Col-0 plants, are listed. Detected peptide numbers by mass spectrometry analysis are shown for each protein.
